# Mapping the impact of microplastics exposure on enteric viral infections in intestinal organoid models

**DOI:** 10.1101/2025.08.19.671065

**Authors:** Jiangrong Zhou, Yilan Zhao, Xingcheng Li, Luc van der Laan, Annemarie C. de Vries, Qiuwei Pan, Pengfei Li

## Abstract

Micro- and nanoplastics (MNPs) are pervasive environmental pollutants increasingly detected in human tissues, yet their long-term health consequences remain poorly understood. The intestinal epithelium, continuously exposed to ingested MNPs and serving as a primary entry site for enteric viruses, provides a critical context to study pollutant-pathogen interactions. We used human intestinal organoids to model chronic exposure, found that prolonged NPs exposure potentially triggered mitochondrial stress and broad metabolic disruption without overt toxicity. This reprogramming altered the host-virus response and also reduced sensitivity to the antiviral treatment. These findings suggest that chronic plastic exposure may subtly but persistently reshape mucosal physiology in ways that alter host-pathogen response and therapeutic efficacy, highlighting an urgent need to address environmental pollutants in infectious disease and public health research.

## Introduction

Plastics are integral to modern life, yet their waste poses a pressing global environmental challenge. Micro- and nanoplastics (MNPs), which originate from the fragmentation of larger plastic items or are intentionally manufactured, are of particular concern, as their non-biodegradable nature enables long-term persistence and progressive accumulation in the environment. They have been detected in marine environments (Zhao et al., 2025), soils (Sajjad et al., 2022,Yoon et al., 2024), air (Abbasi et al., 2019,O’Brien et al., 2023) and also reported to contaminate freshwater (Rodrigues et al., 2018,Zhang et al., 2016,Ajay et al., 2021) and even the human food supply (Kosuth et al., 2018). This ubiquitous presence leads to an inevitable, continuous microplastic exposure for human beings and bioaccumulation across multiple organ systems (Ali et al., 2024,Nihart et al., 2025). Despite recent studies have linked their presence in carotid atheromas to an increased risk of adverse cardiovascular events and found a positive correlation between fecal microplastic levels and inflammatory bowel disease activity (Yan et al., 2022,Marfella et al., 2024), the health consequences of such exposure are not yet well understood.

Another public health concern arises from the potential interaction between MNPs and environmental pathogens. Studies indicate that MNPs can act as vectors for viruses, prolonging their survival (Lu et al., 2022,Zhang et al., 2022), and, in some cases, enhance infectivity (Zhang et al., 2022,Wang et al., 2023). Moreover, in fish models, co-exposure to MPs increased virus mediated mortality compared with viral infection alone (Seeley et al., 2023). Consistently, we observed that nanoparticles enhanced echovirus infection, whereas they interfered with rotavirus infection, suggesting virus-specific interactions (Fig. S1A). Noteworthy, most evidence comes from studies focusing on acute or short-term MNPs exposure, whereas environmental exposure in humans is chronic, raising critical questions about how long-term exposure may affect viral infection.

Ingestion is a major MNPs exposure route for humans (Galloway, 2015), enabling these particles to progressively accumulate in the gut and cause sustained adverse effects such as oxidative stress and inflammation (Hirt and Body-Malapel, 2020). As the intestinal epithelium is also a primary site of enteric viral entry and replication, it offers a relevant setting to investigate how chronic MNPs exposure influences viral infection. Importantly, Polystyrene (PS), a widely used polymer in food containers, contributes substantially to human exposure (Wang et al., 2023,Cassidy and Elyashiv-Barad, 2007). Given its prevalence, PS was selected as a representative microplastic to assess the effects of long-term exposure.

Most studies have used animal models in toxicological assessment, but species differences, limited imaging access, and other constraints reduce their application (Khabib et al., 2022, Ehrenfellner et al., 2017), and traditional cell lines lack the structural and physiological fidelity needed to model authentic host-virus interactions (Speranza, 2023). In contrast, intestinal organoids circumvent these limitations, and preserve the gut’s architecture, cellular diversity, and barrier integrity, while supporting extended culture to assess the chronic toxicity under physiologically relevant conditions.

In this study, we employed this innovative model to pursue three objectives: (i) to establish a chronic exposure system in human intestinal organoids; (ii) to evaluate its impact on viral infection; and (iii) to determine whether it modulates therapy responses.

## Results

### Establishment and characterization of chronic MNPs exposure in human intestinal organoids

To model long-term MNPs exposure in the intestinal epithelium, we established a chronic exposure system using human intestinal organoids (HIOs). Organoids were continuously treated with either 100-nm polystyrene nanoplastics (PS-NPs) or 5-μm microplastics (PS-MPs), with particles replenished at each passage every 7 days (Fig. 1A). During continuous exposure, NPs exhibited pronounced peri-organoid clustering, whereas MPs showed a more diffuse distribution (Fig. S2). Following 4 weeks of exposure, organoids exhibited preserved morphology and viability (Fig. 1B). Confocal microscopy confirmed NPs internalization within organoids, while MPs remained largely extracellular (Fig. 1C). Therefore, based on these findings, NPs were selected for subsequent experiments.

**Figure 1.**
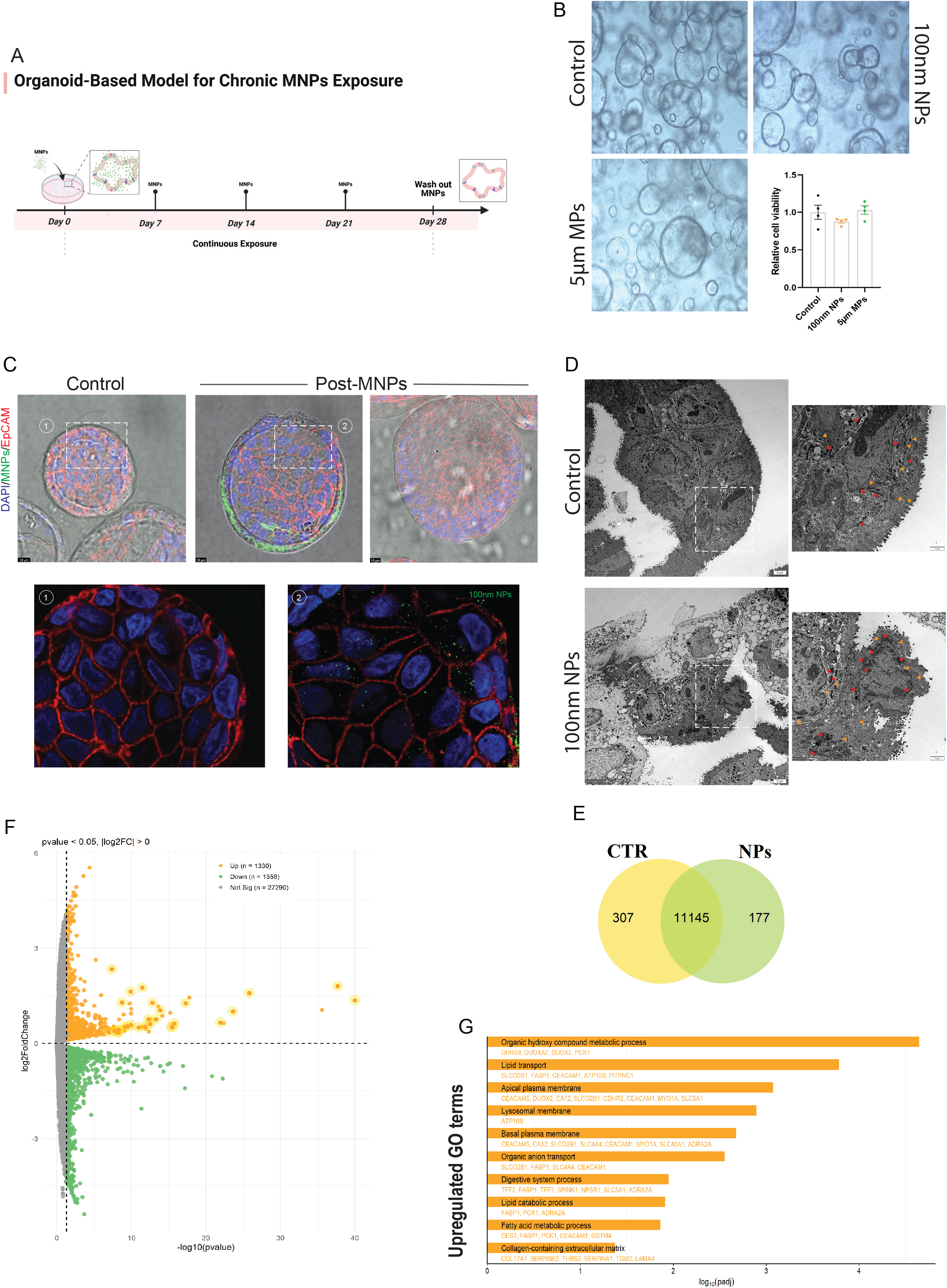
Morphological and transcriptomic effects of chronic PS-MNPs exposure on intestinal organoids. (A) Schematic of the chronic exposure workflow using 100 nm PS-NPs and 5 µm PS-MPs. (B) Immunofluorescence images showing efficient internalization of PS-NPs, with minimal uptake of PS-MPs. Scale bar, 10 μm. (C) Bright-field images demonstrating preserved morphology and relative viability after 4 weeks of MNPs exposure (n = 4). Relative fluorescence units were measured at λ_Exc 530 nm / λ_Em 590 nm. (D) Representative TEM images. Control organoids show intact epithelial ultrastructure with normal mitochondria (red arrowheads) and rough endoplasmic reticulum (RER; orange arrowheads), whereas NPs-exposed organoids display reduced cytoplasmic density and abnormal mitochondria and RER. (E) Venn diagram showing 177 differentially expressed genes (DEGs) unique to NPs-exposed organoids and 307 unique to controls. (F) Volcano plot of differentially expressed genes (DEGs) in NPs-exposed versus control organoids, highlighting significantly upregulated genes (CEACAM5, DHRS9, REG4, DUOXA2, DUOX2, MMP1, COL17A1, SLC2A3) (p < 0.05). (G) Top 10 significantly enriched pathways identified by Gene Ontology (GO) analysis of NPs-exposed organoids compared with controls.

Transmission electron microscopy (TEM) revealed ultrastructural differences between control and NPs-exposed organoids (Fig. 1D). Control organoids possessed intact membranes, moderate cytoplasmic electron density, mitochondria (red arrowheads) with normal oval-to-elongated morphology, and rough endoplasmic reticulum (RER, orange arrowheads) with intact cisternal architecture. In contrast, NPs-exposed organoids exhibited reduced cytoplasmic electron density, pronounced mitochondrial cristae dilation with small vacuoles, and mild RER cisternal widening accompanied by partial ribosome loss.

Transcriptomic profiling further corroborated the morphological alterations observed. The Venn diagram (Fig. 1E) demonstrated 177 DEGs unique to NPs-treated organoids and 307 unique to controls, with 11,145 shared genes, while principal component analysis (Fig. S3) demonstrated clear separation between NPs-treated and control organoids, indicating robust transcriptomic divergence. Volcano plot analysis (Fig. 1F) revealed a balanced transcriptional response, with 1,330 genes significantly upregulated and 1,358 downregulated (p < 0.05) in NPs-treated versus control organoids; prominently upregulated genes (CEACAM5, DHRS9, REG4, DUOXA2, DUOX2, MMP1, COL17A1, SLC2A3) are associated with epithelial barrier integrity, oxidative stress responses, and metabolic regulation. GO enrichment (Fig. 1G) highlighted major shifts in metabolic and transport-related pathways, including lipid metabolism, membrane restructuring, transport regulation, and digestive activity, alongside ECM remodeling, suggesting that chronic NPs exposure drives broad metabolic reprogramming, altered transport capacity, and potential disruption of epithelial functional homeostasis.

Together, chronic NPs exposure in HIO revealed persistent particle–epithelium interaction, active internalization, and early ultrastructural changes without compromising viability, while triggering broad metabolic reprogramming and functional shifts in epithelial homeostasis, further providing a robust platform to study MNPs impacts on viral infection and host responses.

### Chronic NPs exposure affects enteric virus infection and reprograms the host-virus transcriptional response

To investigate how chronic NPs exposure influences viral infection, we first examined echovirus 1 (EV1) (Fig. 2A), an enteric virus transmitted via the fecal-oral route that infects the intestinal epithelium early in its life cycle (Wells et al., 2022). Chronic NPs exposure markedly reduced EV1 infection, as demonstrated by qRT-PCR quantification of viral RNA (Fig. 2B) and TCID_50_ assays showing significantly lower infectious titers in NPs-treated organoids (Fig. 2C). Immunofluorescence staining corroborated these findings, with abundant dsRNA signal observed in control organoids, consistent with robust replication, whereas NPs-exposed organoids exhibited weaker, focal dsRNA staining that appeared to colocalize with NPs (Fig. 2D). A second enteric virus, rotavirus (RV), was then assessed. Similar to EV1, RV infection was significantly reduced in NPs-treated organoids (Fig. 2E-F), and confocal imaging revealed diminished dsRNA signal (Fig. S4A). In addition, NPs exposure attenuated virus-induced cytopathic effects (Fig. 2G; Fig. S4B-C). Collectively, these results indicate that chronic NPs exposure alters both infection and cytopathic effects of single- and double-stranded RNA enteric viruses in HIO.

**Figure 2.**
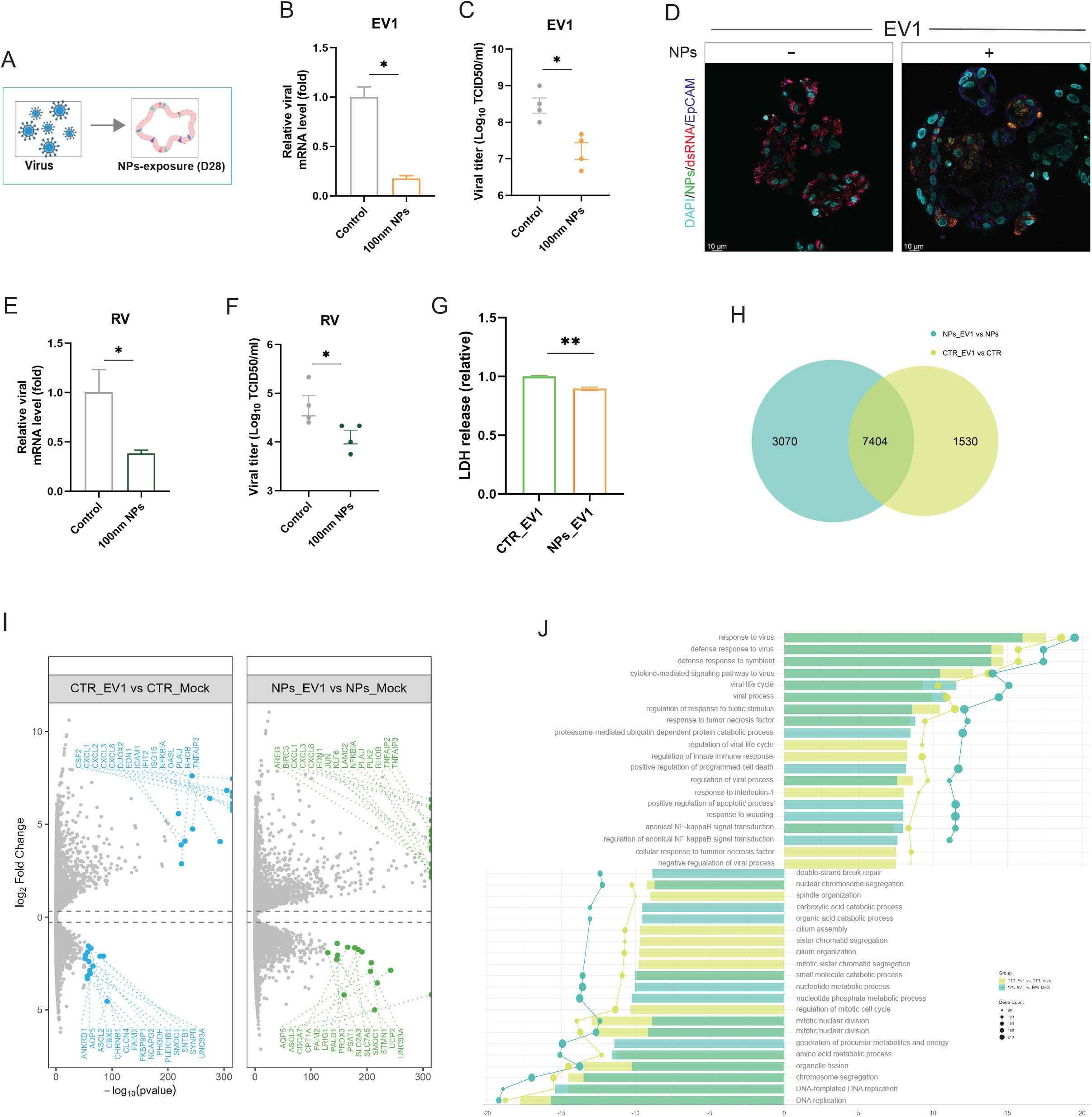
Chronic NPs exposure remodels the host–virus response. (A) Schematic of the experimental design for chronic NPs exposure followed by viral infection. (B-C) qRT-PCR quantification of (echovirus 1) EV1 RNA levels (B) and infectious viral titers (C) in control and NPs-exposed organoids. (D) Confocal images of replicating viral dsRNA (red) in EV1-infected organoids at 48 h post-infection, with fluorescently labeled nanoparticles (green) in NPs-exposed and control conditions. Scale bar, 10 μm. (E-F) qRT-PCR quantification of rotavirus (RV) RNA levels (E) and infectious viral titers (F) in control and NPs-exposed organoids. (G) Quantification of lactate dehydrogenase (LDH) release at 48 h post-inoculation in EV1-infected control and NPs-exposed organoids (n = 4). (H) Venn diagram of unique DEGs in EV1-infected NPs-exposed and control organoids relative to uninfected organoids at 48 h post-inoculation (n = 4). (J) Volcano plot of significantly upregulated and downregulated DEGs in EV1-infected NPs-exposed and control organoids compared with their respective uninfected organoids at 48 h post-inoculation (p < 0.05). (K)Top 15 significantly enriched Gene Ontology (GO) terms in EV1-infected NPs-exposed organoids versus uninfected organoids, and in infected control organoids relative to uninfected organoids at 48 h post-inoculation (n = 4).

To explore the underlying host-virus response in organoids with or without prior NPs exposure, we selected EV1 as a model virus and performed genome-wide transcriptomic analysis. Intriguingly, NPs exposed groups showed an altered host transcriptional response to EV1 (Fig. 2H). Following EV1 infection, both the control and NPs-exposed organoids activated inflammatory associated genes like CXCL1, CXCL3, and PLAU, indicating preserved innate viral response. In controls, EV1 triggered strong induction of canonical antiviral genes IFIT2 and ISG15, along with immune cell recruitment factors, and enrichment of pathways involved in viral process regulation and NF-κB modulation, accompanied by broad suppression of metabolic and biosynthetic programs (Fig. 2I-J). NPs-exposed organoids instead showed enrichment of apoptotic signaling, programmed cell death and proteasome-mediated degradation, coupled with pronounced downregulation of energy metabolism and precursor metabolite generation (Fig. 2J). These findings indicate that NPs exposure reprograms the host-virus response from a coordinated antiviral-metabolic suppression to a compromised antiviral state with enhanced cell death responses and impaired bioenergetic synthesis.

### Chronic NPs exposure attenuates antiviral drug responsiveness

Since NPs-treated organoids exhibit a distinct host-virus transcriptional response, we next investigated whether antiviral efficacy was affected. EV1-infected organoids, with or without prior NPs exposure, were treated with pleconaril, a capsid-binding inhibitor. RT-qPCR quantification of viral RNA revealed a clear dose-dependent inhibition of EV1 infection in both groups (Fig. S5A-B). However, NPs-exposed organoids consistently retained higher residual viral RNA levels than matched controls, indicating reduced pleconaril efficacy (Fig. 3A).

**Figure 3.**
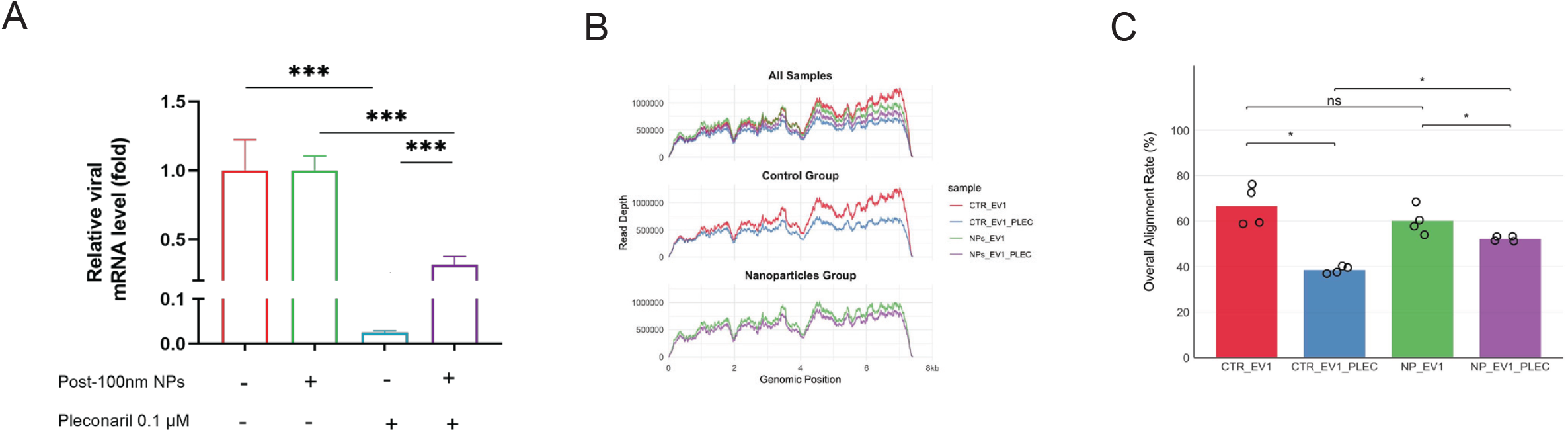
Chronic NPs exposure alters antiviral drug responsiveness in EV1-infected organoids. (A) Quantification of viral RNA levels in EV1-infected NPs-exposed and control organoids with or without 0.1 µM pleconaril treatment for 48 h (n = 8). (B-C) Whole-genome view of mapped transcripts by RNA-seq across the EV1 genome and overall alignment rates of viral reads to the EV1 genome in infected NPs-exposed and control organoids, with or without pleconaril treatment (n = 4).

Genome-wide viral RNA analysis reinforced these findings. RNA sequencing revealed that in control organoids, pleconaril significantly reduced EV1 reads depth across the genome, whereas NPs-exposed organoids showed only modest reductions, with coverage profiles for treated and untreated samples remaining closely aligned (Fig. 3B). Consistently, the overall alignment rate, the proportion of sequencing reads mapping to the EV1 genome, decreased after pleconaril treatment in both groups but remained substantially higher in NPs-treated organoids (Fig. 3C).

Together, these results indicate that while pleconaril remains active against EV1, chronic NPs exposure compromises its efficacy. This reduced responsiveness likely reflects NPs-induced reprogramming of host-virus-drug interactions.

## Discussion

Organoid-based models have recently emerged as powerful tools for studying MNPs toxicity, providing insights into organ-specific vulnerabilities and mechanistic pathways that traditional in vitro and animal models often fail to capture(Park et al., 2023,Hou et al., 2022,Cheng et al., 2024,Cheng et al., 2022,Cheng et al., 2023). Moreover, Park et al. demonstrated that MNPs-induced toxic effects in human colon organoids more closely resembled in vivo responses than those observed in conventional colon cell cultures(Park, et al., 2023). Nevertheless, most studies to date have focused on short-term or acute exposures, leaving the long-term health risks associated with chronic MNPs exposure largely unexplored.

In this study, we applied human intestinal organoids to investigate the effects of prolonged MNPs exposure, with the gut epithelium serving as a physiologically relevant target site of continuous contact with environmental pollutants. However, organoids cultured in Matrigel retain a 3D structure but exhibit reversed polarity, with the basal surface exposed outward, unlike the native intestine where the apical side faces the lumen (Lee et al., 2024). While dynamic exposure systems such as rotating-flask agitation can invert polarity, and indeed facilitate apical contact (Cheng, et al., 2022,Cheng, et al., 2023), these approaches require specialized equipment and remain technically demanding.

Here, we developed an organoid fragmentation-MNPs coculture model that enabled both the outer and inner cell surfaces to directly interact with MNPs. Importantly, the fragmented organoid-MNPs mixture could be reseeded and serially passaged under repeated exposure, enabling long-term assessment over 28 days, significantly longer than the 48 h or 14-day exposures previously reported (Park, et al., 2023,Hou, et al., 2022), while maintaining organoid morphology and viability. Our results showed that nano-sized plastics were readily internalized by organoid cells, whereas larger microplastics remained predominantly extracellular, confirming size-dependent uptake feature consistent with previous observations (Lu et al., 2022,Shen et al., 2022,Chen et al., 2020).

Although we did not directly trace the subcellular localization of particles, several lines of evidence point to mitochondrial involvement. PS-NPs exposure induced cristae dilation, vacuolization, and RER alterations, ultrastructural changes consistent with mitochondrial stress and functional disruption. These observations align with prior reports (Qin et al., 2022,Jeong et al., 2016), and are further supported by our transcriptomic data, which revealed broad reprogramming of metabolic and transport pathways together with signatures of oxidative stress and epithelial remodeling. Notably, similar metabolic disruptions, including ROS accumulation and loss of mitochondrial potential, have been also observed in mouse intestinal organoids (Xuan et al., 2024), highlighting mitochondria as a conserved point of vulnerability to MNPs exposure across species.

The intestinal epithelium represents the primary barrier of host defense against pathogens, and its integrity depends on balanced cellular function. Our results suggest that chronic MNPs exposure may compromise this barrier, potentially weakening mucosal protection. Beyond characterizing PS-NPs accumulation and cellular impacts, we also examined potential effects on viral infection. Interestingly, NPs-treated organoids showed reduced viral RNA levels, lower infectious titers, and attenuated dsRNA staining compared with controls. These outcomes contrast with previous reports describing microplastics as enhancers of viral infectivity and virus-mediated mortality (Zhang, et al., 2022,Wang, et al., 2023,Seeley, et al., 2023), suggesting that exposure duration, particle size, and experimental context critically shape biological outcomes.

To elucidate the underlying mechanism, the most relevant finding was the distinct transcriptomic host-virus responses in NPs treated versus control organoids. Control organoids mounted a more complete antiviral program, characterized by robust interferon signaling and activation of canonical antiviral pathways. In contrast, NPs-exposed organoids exhibited attenuated interferon responses and weaker activation of these pathways. Surprisingly, despite this blunted antiviral profile, NPs-exposed organoids showed reduced viral replication. This paradox may be possibly explained by chronic NPs exposure inducing metabolic disturbance and oxidative stress, thereby dramatically creating an epithelial environment less permissive to viral propagation (Kleinehr et al., 2025,Khan et al., 2021,Palmer, 2022), but further insights are needed. Importantly, while this effect might superficially appear protective against certain pathogens, it is in fact particularly alarming, as our model demonstrates clear potential adverse effects as described above.

Apart from altering host-virus interactions, chronic NPs exposure may also compromise antiviral drug efficacy. Although pleconaril, a capsid-binding antiviral, retained dose-dependent inhibitory activity against EV1 in both NPs-exposed and control organoids, its maximal efficacy was diminished following chronic NPs exposure. This attenuation was clearly confirmed in genome-wide analyses of viral abundance and integrity. While the precise mechanisms remain unclear, they may involve NPs-driven modulation of host pathways required for drug activity, warranting further investigation.

From a public health perspective, our study raises the possibility that chronic ingestion of micro- and nanoplastics could subtly but persistently alter mucosal function, potentially affecting both susceptibility to enteric pathogens and responses to antiviral therapy. Future studies should validate these findings in vivo, extend them to additional pathogens, and define the molecular links between MNPs-induced metabolic rewiring, altered host-virus interactions, and pharmacological responsiveness.

These findings also underscore the value of intestinal organoids as powerful tools for assessing the health impacts of environmental pollutants. By applying a chronic exposure paradigm, we move beyond the short-term or acute models that dominate the field and provide a multidimensional perspective that links chronic MNPs exposure to cellular function remodeling, infectious disease outcomes, and therapeutic efficacy. A particular strength of our model lies in its simple and adaptable design, which not only makes it feasible to investigate the long-term toxicity of MNPs but also offers a broadly applicable platform for evaluating the effects of other environmental pollutants.

This study has several limitations. First, although intestinal organoids recapitulate key aspects of epithelial physiology, they lack systemic immune components, and the influence of the gut microbiome, all of which are critical modulators of host responses to both pollutants and pathogens. Second, we focused on a single type of polystyrene particle, and future work should evaluate whether the observed effects extend to other plastic compositions, particle sizes, and shapes more representative of real-world exposures. Third, while our chronic exposure model provides valuable insight into long-term epithelial responses, the dosing paradigm may not fully capture the fluctuating and heterogeneous exposures experienced in vivo. Finally, although we observed ultrastructural and transcriptomic changes suggestive of metabolic and antiviral reprogramming, functional validation of the implicated pathways will be necessary to establish causality. These limitations also highlight key avenues for future research, including incorporating immune and microbial components into organoid systems, expanding to diverse environmental pollutants, and mechanistically dissecting the molecular drivers of pollutant-host-pathogen interactions.

In summary, our findings demonstrate that chronic MNPs exposure reprograms intestinal epithelial physiology in ways that alter host-virus interactions and diminish antiviral drug efficacy. While these effects may superficially appear protective against infection, they instead reveal an underrecognized risk of pollutant-driven trade-offs in mucosal defense and therapeutic responsiveness. Together, this work highlights the importance of considering chronic environmental pollutant exposure in infection biology and underscores the utility of intestinal organoids as scalable, physiologically relevant platforms for assessing the health impacts of particulate pollutants.

## Materials and methods

### Polystyrene micro(nano)plastics (PS-MNPs)

Fluoresbrite^®^ YG Carboxylate Microspheres (0.10 μm; Polysciences, Cat. No. 16662-10, Tebu-Bio BV), polystyrene microparticles (5 μm; Merck Life Science NV, Cat. No. 79633-10ML-F), and DiagPoly™ Carboxyl Fluorescent Polystyrene Particles, Green (5 μm; CD Bioparticles, Cat. No. DCFG-L011) were used in this study. The stock solutions were diluted in organoid expansion medium to prepare polystyrene (PS) culture solutions before experiment.

### Intestinal organoids culture and ethics

Human intestinal organoids (HIO) were established as described previously (Bijvelds et al., 2022,Yin et al., 2015). Organoids were cultured in organoid expansion medium (OEM) based on Advanced DMEM/F12 (Invitrogen), supplemented with 1% penicillin/streptomycin (Life Technologies), 10 mM HEPES, 1× GlutaMAX, 1 mM N2, 1 mM B27 (all from Invitrogen), 1 μM N-acetylcysteine (Sigma), and the following growth factors: 50 ng/L mouse epidermal growth factor (mEGF), 50% Wnt3a-conditioned medium (WCM), 10% noggin-conditioned medium (NCM), 20% Rspo1-conditioned medium, 10 μM nicotinamide (Sigma), 10 nM gastrin (Sigma), 500 nM A83–01 (Tocris), and 10 μM SB202190 (Sigma). The medium was refreshed every 2-3 days, and organoids were passaged at a 1:4 ratio every 6-7 days. Use of human intestinal tissue was approved by the Medical Ethical Council of Erasmus MC with informed consent (MEC-2021-0432; MEC-2023-0629).

### Cell lines and viruses

Adenocarcinomic human alveolar basal epithelial cells (A549) and African green monkey kidney cells (MA-104) were cultured in Dulbecco’s modified Eagle medium (DMEM; Lonza Biowhittaker) supplemented with 10% (v/v) heat-inactivated fetal calf serum (FCS; Sigma–Aldrich), 100 IU/mL penicillin, and 100 μg/mL streptomycin (Gibco). Echovirus 1 (EV1; GenBank: AF029859) and echovirus 6 (EV6; GenBank: JQ929657) were propagated in A549 cells, while rotavirus strain SA11 (GenBank: X16830) was propagated in MA-104 cells. All cell lines were authenticated by genotyping and confirmed to be mycoplasma-free.

### Establishment and Characterization of MNPs-Exposed Organoids

Human intestinal organoids were cultured in Matrigel until >75% confluence (diameter ≈ 200 µm). For PS-MNPs exposure, organoids were washed with cold Advanced DMEM/F12, centrifuged (300 × g, 5 min), and mechanically dissociated. Pellets were resuspended in expansion medium containing PS-MNPs (exposure concentration: 300 µg/mL; experimental) or PS-MNPs free medium (control), mixed with Matrigel (2:8, v/v) by pipetting ×5, and seeded into plates. After Matrigel polymerization, cultures were overlaid with expansion medium (refreshed every 2 days). Organoids were passaged every 6-7 days, with repeated MNPs treatment at the same concentration for experimental wells. The concentration of 300 µg/mL was selected based on prior in vitro exposure studies showing detectable microplastic uptake without overt cytotoxicity (Koner et al., 2025), enabling chronic exposure modeling. After exposure, the organoids are collected for following experiments, as described below. LEICA SP8 confocal microscopy were used for visualization and transmission electron microscopy (TEM) to examine ultrastructural changes, confirm MNPs internalization and determine subcellular localization.

### Virus inoculation

Organoids were mechanically fragmented and exposed to viral particles for 2 h at 37 °C with rotavirus, EV1, or EV6. Each organoid preparation was inoculated with approximately 4 × 10^4 PFU of rotavirus (10^3.8 PFU) or 2 × 10^4 PFU of echovirus; rotavirus was pre-activated with 0.05% trypsin for 10 min at 37 °C. To enhance infection efficiency, the organoid-virus mixture was gently resuspended every 30 min during the inoculation period. Subsequently, the organoids were centrifuged at 300 × g for 5 min at 4 °C, the supernatant was discarded, and the pellets were washed three times with Advanced DMEM/F12 to remove residual viral particles. Following infection, organoids were seeded into wells, with non-infected organoids processed under identical conditions serving as mock controls.

### Immunofluorescence staining and confocal imaging

Organoids were fixed with 4% paraformaldehyde (PFA) for 15 min. Then the samples were gently rinsed 3 times with PBS, followed by permeabilizing with PBS containing 0.2% (vol/vol) Triton X-100 for 10⍰min. Next, samples were twice rinsed with PBS for 5⍰min, followed by incubation with blocking solution (5% donkey serum, 1% bovine serum albumin, 0.2% Triton X-100 in PBS) at room temperature for 1⍰hour. Primary antibodies used in this study are as follows: anti-EpCAM (1:500, rabbit), anti-dsRNA antibody (1:500, mouse). Sample were then washed 3 times for 5⍰min each in PBS prior to 1⍰hour incubation with 1:1000 dilutions of the secondary antibodies including anti-mouse IgG (H+L, Alexa Fluor^®^ 594) and the anti-rabbit IgG (H+L, Alexa Fluor^®^ 647). Nuclei were stained with DAPI (4, 6-diamidino-2-phenylindole; Invitrogen). At last, stained samples were visualized using a Leica SP5 confocal microscope with a 40× oil immersion objective to analyze the stained cellular structures.

### QRT-PCR quantification of gene expression

Total RNA was extracted using the Macherey-Nagel NucleoSpin RNA II Kit (Bioke, Leiden, Netherlands). The concentration and purity were measured using the Nanodrop ND-1000 (Wilmington, DE, USA). Gene expression levels were quantified by SYBR Green–based qRT-PCR using the Applied Borganoidsystems SYBR Green PCR Master Mix (Thermo Fisher Scientific Life Sciences) with the StepOnePlus System (Thermo Fisher Scientific Life Sciences). The housekeeping gene used for normalization was glyceraldehyde 3-phosphate dehydrogenase (GAPDH). Relative gene expression was normalized to GAPDH using the 2^−ΔΔCT method (ΔΔCT = ΔCT_sample − ΔCT_control). Each qRT-PCR experiment included template control and reverse transcriptase control.

### Cell Viability and Cytotoxicity Assessment

Organoids were incubated with AlamarBlue reagent (1:20; Invitrogen, DAL1100) in culture medium for 2 h at 37 °C. Subsequently, 100 μL of medium from each well was collected to assess metabolic activity, with each sample measured in duplicate. Fluorescence was recorded using a CytoFluor Series 4000 plate reader (PerSeptive Biosystems; 530/25 nm excitation, 590/35 nm emission).

Cytotoxicity was quantified by lactate dehydrogenase (LDH) release using the CytoTox 96^®^ Non-Radioactive Cytotoxicity Assay (Promega, G1780) per manufacturer’s instructions. Supernatants from virus-infected organoids were analyzed at 490 nm, and LDH release was expressed as a percentage of total LDH after complete lysis.

### Genome-wide RNA sequencing and data analysis

Organoids with or without PS-NPs exposure were infected with EV1 as described above and harvested at 48 h post-infection. For antiviral treatment, infected organoids were exposed to 0.1 μM pleconaril starting 1 h post-infection and maintained until 48 h post-infection. In parallel, uninfected organoids were cultured under identical conditions as negative controls. Total RNA was extracted using the Macherey-Nagel NucleoSpin RNA II Kit (Bioke, Netherlands) and quantified with the Bioanalyzer RNA 6000 Picochip. RNA sequencing was performed by Novogene using a paired-end 150 bp (PE150) strategy to assess transcriptional changes associated with chronic PS-NPs exposure and EV1 infection.

### Statistics

Statistical analyses were performed using GraphPad Prism 8 (GraphPad Software, San Diego, USA). Data are presented as the mean⍰±⍰standard error of the mean (s.e.m.). Comparisons between two groups were made using an unpaired two-tailed Student’s t-test. Asterisks indicated the degree of significant differences compared with the controls (^*^, p < 0.05; ^**^, p < 0.01; ^**^, p < 0.001; ^****^, p < 0.0001).

## Author contributions

J.R.Z., Q.W.P., and P.F.L. conceived the study. J.R.Z., Y.L.Z., and X.C.L. designed the experiments. J.R.Z. and Y.L.Z. performed data analysis and visualization. J.R.Z. and X.C.L. cultured intestinal organoids. Y.L.Z. propagated echovirus and rotavirus. J.R.Z. and P.F.L. wrote and revised the manuscript. L.L., Q.W.P., and A.C.V. critically reviewed the manuscript. P.F.L. supervised the project and secured funding.

