## Supplementary Figures 1-5 for "Mapping the impact of microplastics exposure on enteric viral infections in intestinal organoid models"

**Supplementary Figure 1**

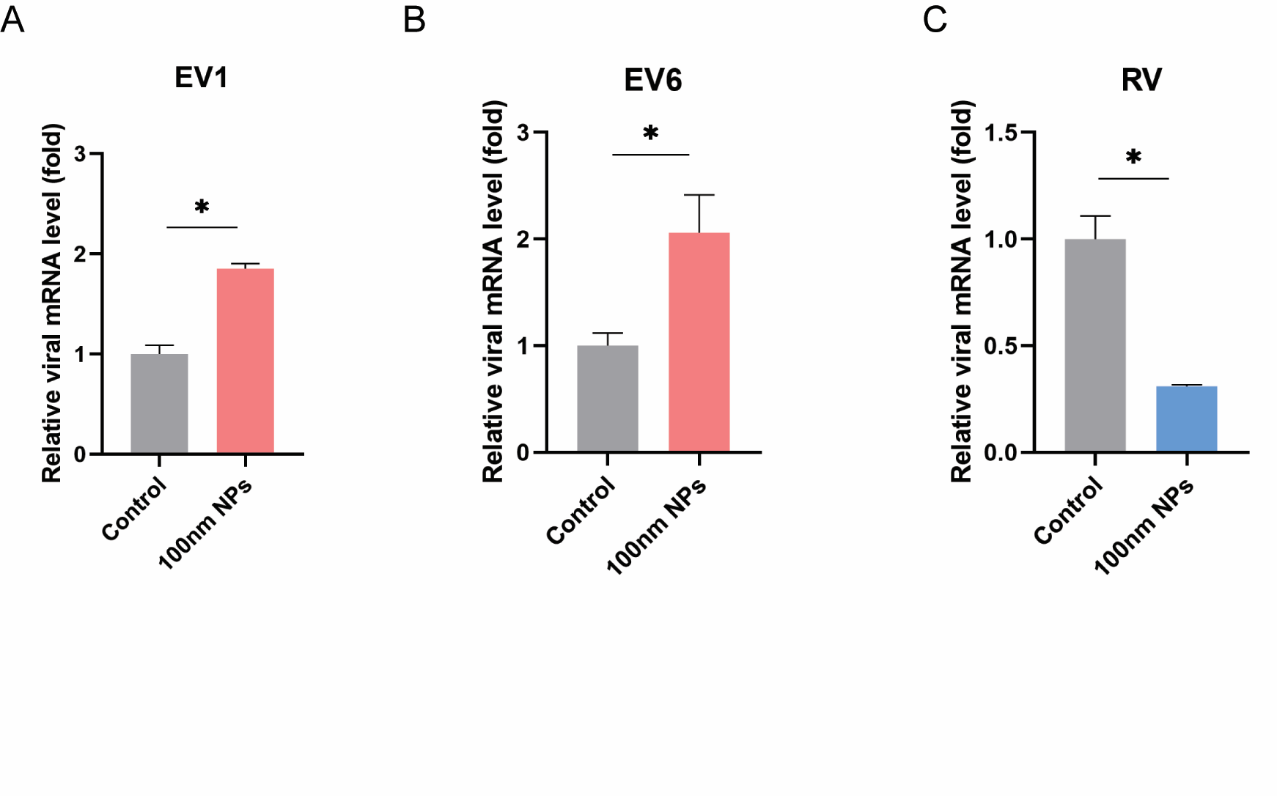

**Fig. 1** (A–C) Quantification of viral RNA expression following infection with echovirus 1 (EV1) (A), echovirus 6 (EV6) (B), and rotavirus (RV) (C) in organoids co-exposed to nanoparticles (NPs), with infections without NP exposure serving as controls (n = 4).

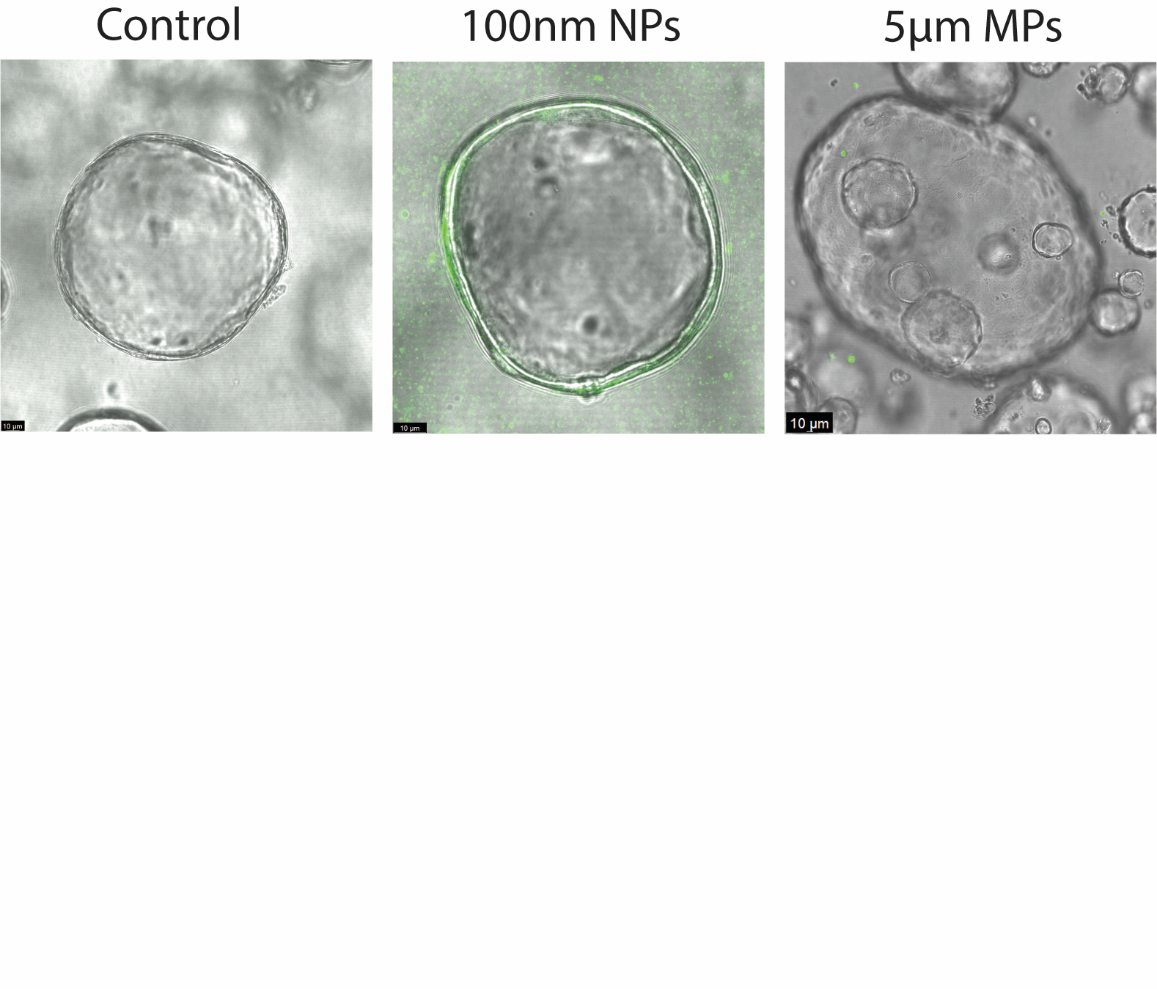
**Supplementary Figure 2**

**Fig. 2** Confocal image of organoids co-cultured with 100-nm polystyrene nanoplastics (PS-NPs) or 5-μm microplastics (PS-MPs) (green), respectively. Scale bar, 10 μm.

**Supplementary Figure 3**

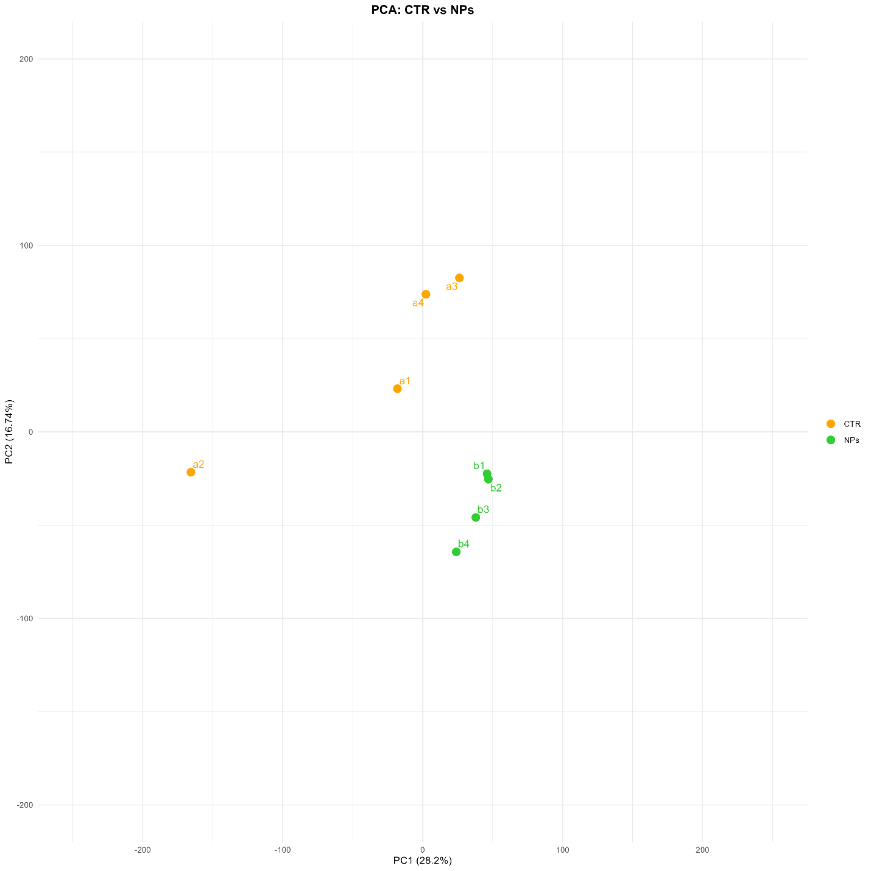

**Fig. 3** Principal component analysis of NPs-exposed and control organoids (n=4)

**Supplementary Figure 4**

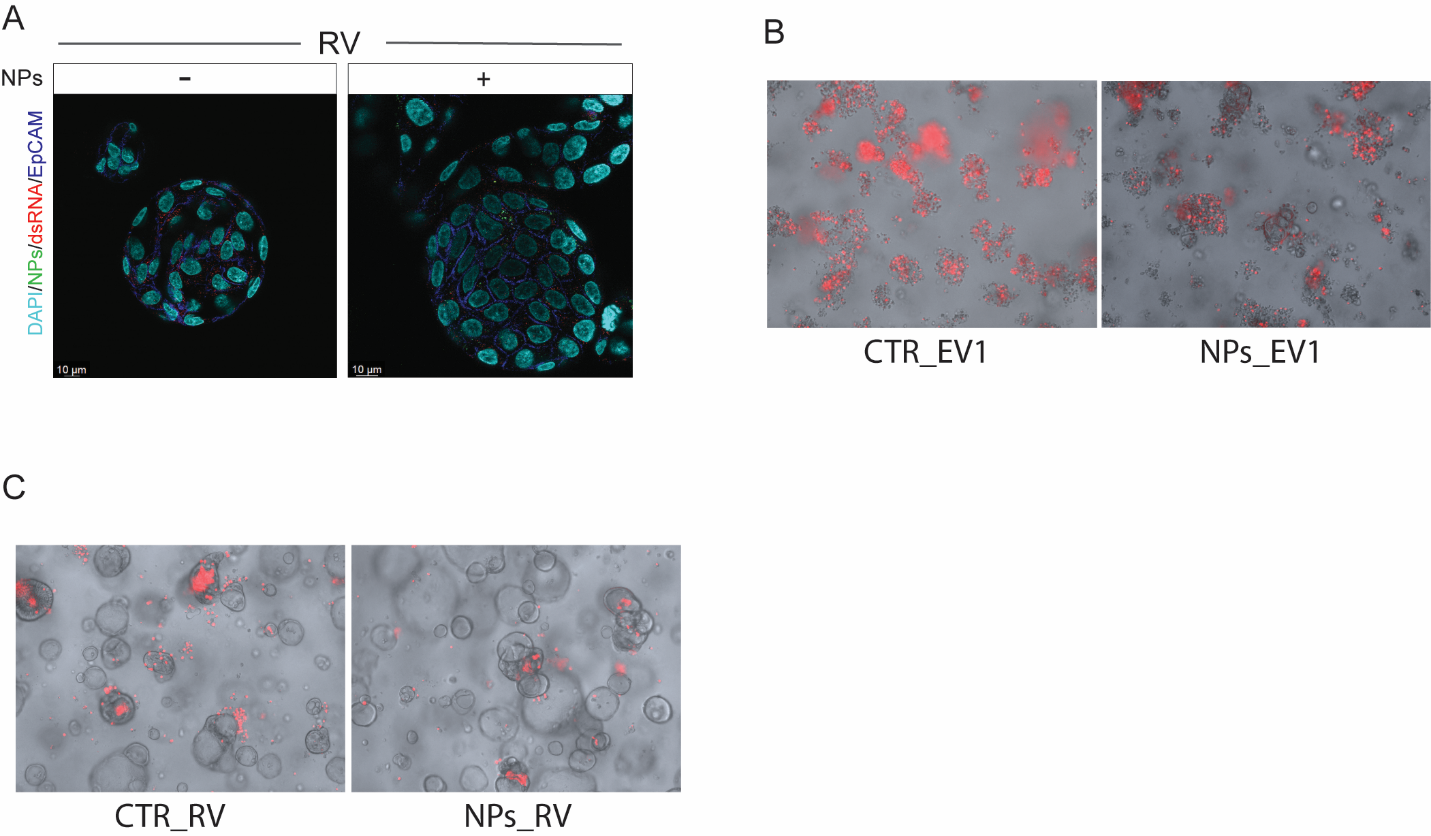

**Fig. 4** (A) Confocal images of replicating viral dsRNA (red) in RV-infected organoids at 48 h post-infection, with fluorescently labeled nanoparticles (green) in NPs-exposed and control conditions. Scale bar, 10 μm. (B-C) Propidium iodide (PI) staining showing EV1- (B) and RV- (C) induced cell death in NPs-exposed and control organoids. Images were acquired at 10× magnification.

**Supplementary Figure 5**

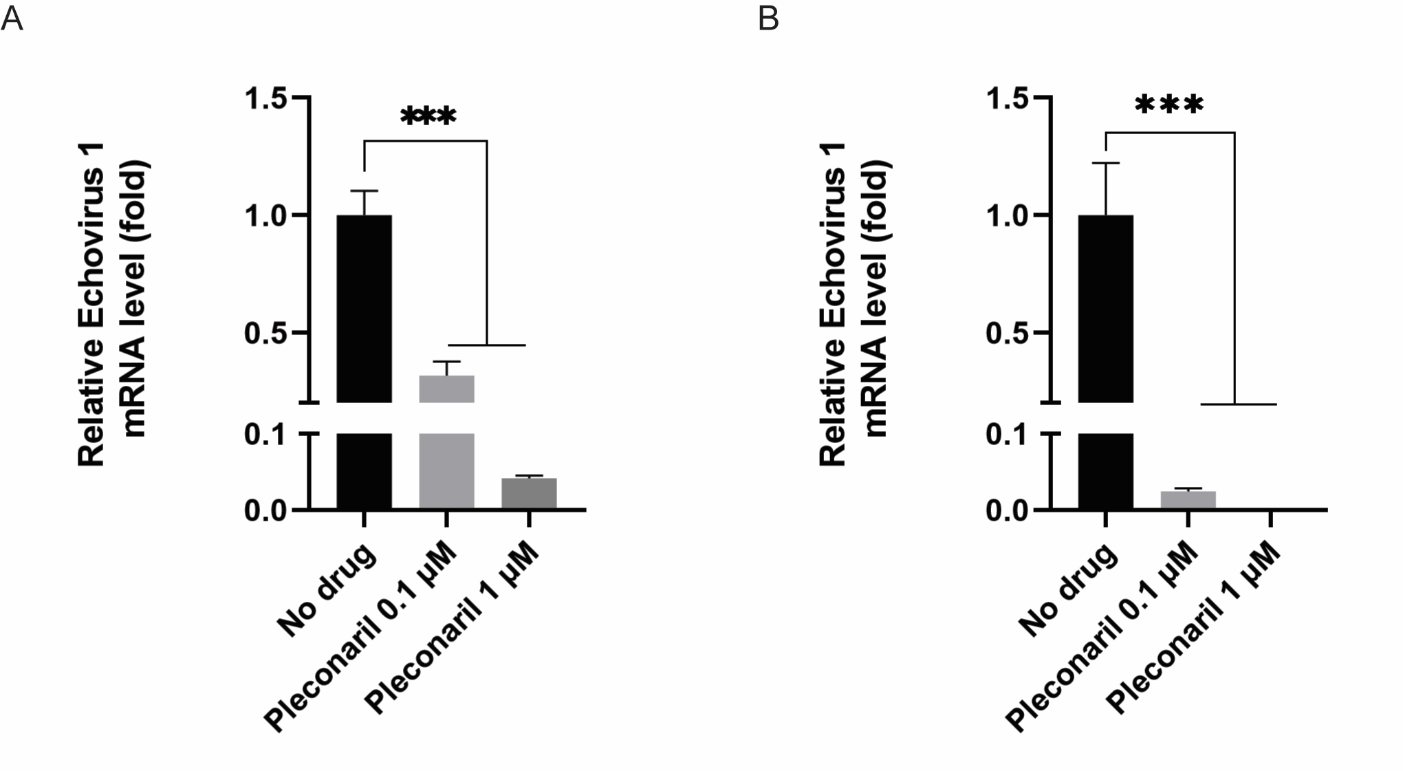

**Fig. 5** (A-B) Quantification of viral RNA levels in EV1-infected organoids exposed to nanoparticles (A) or in control organoids (B) treated with pleconaril (0.1 µM or 1 µM) for 48 h. Data are shown relative to infected organoids without treatment (n =8).
